# ReachingBot: an automated and scalable benchtop device for highly parallel Single Pellet Reach-and-Grasp training and assessment in mice

**DOI:** 10.1101/2022.06.17.496542

**Authors:** Sotiris G. Kakanos, Dhireshan Gadiagellan, Eugene Kim, Diana Cash, Lawrence D. F. Moon

## Abstract

The single pellet reaching and grasp (SPRG) task is a behavioural assay widely used to study motor learning, control and recovery after nervous system injury in animals. The manual training and assessment of the SPRG is labour intensive and time consuming and has led to the development of multiple devices which automate the SPRG task. Current state-of-the-art desktop methods either still require attendance, manual classification of trial outcome, or expensive locally-installed hardware such as graphical processing units (GPUs). Here, using robotics, computer vision, and machine learning analysis of videos, we describe a novel cost-effective benchtop device that can be left unattended, presents pellets to mice automatically, video records each trial, and, using two supervised learning algorithms, classifies the outcome of each trial automatically with an accuracy of greater than 94% without the use of GPUs. Finally, the device is simple in design with few components meaning manufacturing at scale is straightforward and, with few moving parts, reliable and robust. Our devices can also be operated using our cross-platform Graphical User Interface (GUI), meaning no knowledge of programming is required by its users.

We show that these devices can train 30 mice with them collectively performing ~83,000 trials over 3 months, saving users an estimated 8 and half hours of labour per day. Over five weeks, most mice undertook more trials per session and retrieved more pellets successfully. 21 out of 30 mice retrieved at least 40% of pellets successfully in at least one session during the training period. Devices measured motor deficits induced in mice by a focal ischaemic stroke; some mice showed large persistent deficits whilst others showed only transient deficits. This highlights the heterogeneity in reaching outcomes following stroke. We conjecture that reach-and-grasp is represented in motor cortex bilaterally but with greater asymmetry in some mice than in others. We predict that bilateral lesions of motor cortex would cause long-lasting deficits in reach-and-grasp in mice.

We propose a strategy for preclinical evaluation of novel therapeutics that improve reach-and-grasp by pre-screening a large cohort of mice automatically and excluding those that fail to achieve pre-specific success rates, which generates a cohort of mice trained with consistent performance levels, suitable for randomization to treatment arms in a preclinical study. Well-powered sample sizes are easily achievable. Highly parallel automated training and assessment should accelerate the development of new therapies for movement disorders.

## Introduction

The single pellet reaching and grasping (SPRG) task was first described in 1990 by Whishaw and Pellis for use in rodents (Whishaw and Pellis, 1990). It is used to mimic reaching and grasping in humans, to study both motor learning (Mykins et al., 2021) and recovery after brain injury preclinically (Wahl et al., 2014). The SPRG task challenges animals to reach for a food reward through a thin vertical window from within a clear acrylic box. Initially, animals first exposed to the task fail to retrieve pellets, but with training their success rate increases. The ratio of successful grasps compared to failed grasps of the food pellet reward is often used as a principle behavioural outcome measure: the rate at which this ratio increases can be used in motor learning studies, (Mykins et al., 2021, Bova et al., 2021) or in assessing deficits and recovery in animal models of stroke (Wahl et al., 2014), spinal cord injury (Fenrich et al., 2016), Huntington’s Disease (Glangetas et al., 2020) or Parkinson’s Disease (Bova et al., 2020). Other measurements can also be taken from animals executing this task, such as the kinematics of the reaching trajectory, the velocity of the reach and the number of reach attempts, for example (Bova et al., 2021, Mykins et al., 2021).

Researchers have traditionally trained and assessed rodents in SPRG manually: by placing pellets to be grasped by rodents by hand and then by manually scoring successful and failed retrievals. Manual training and assessment is time consuming: each SPRG session typically lasts 20 minutes which requires the full attention of the experimenter so that animal performance can be scored and pellets replaced. Manual scoring also limits the number of metrics one can gather from each reach, requiring video recording of trials for further analysis “offline”. For this reason, various research groups have begun automating this task as well as other assays for the interrogation of motor function in animals reviewed thoroughly elsewhere (Mah et al., 2021).

Overall there are two broad approaches to tackle the automation of SPRG: (1) using in-cage devices, allowing animals to execute the task in their home-cage environment (Salameh et al., 2020, Kakanos, 2020), or (2) by using a benchtop solution, where the task is done in an apparatus outside of the home-cage (Fenrich et al., 2016, Torres-Espín et al., 2018, Bowles et al., 2021). Both solutions have advantages and disadvantages. One advantage of in-cage devices is that rodents can participate freely 24 hours a day. Moreover, rodents can be socially housed and assessed independently if they are implanted subcutaneously with radiofrequency identification (RFID) tags that the device is designed to read (Kakanos, 2020). Further, any stress induced from moving animals to and from their home cages to testing rooms is removed, and the set up requires little to no technician time. However, in-cage devices must function in low to no light conditions during their dark cycle, must be durable in-cage and operate reliably (e.g., without jamming due to bedding and wood chips typically used in animal cages), and, if used in cages with multiple animals, devices must discern which animal executes the task and might even tailor the task parameters (e.g., location of pellet) to the individual animal. Cages might also need to be modified in order to accommodate the hardware, and having robotic devices within the cages of rodents poses safety risks, and limits the space available to the animals. Further, having one device per cage of animals dilutes the amount of exposure each animal has to the task; our data (unpublished) suggest that the degree of participation among group housed mice varies significantly. All of the above requirements add layers of complexity to the in-cage hardware solution.

Benchtop devices have the advantage over those that are in-cage because the device needs to solve fewer problems. For example, it does not need to operate in the dark, it does not need to discern which animal uses the device and it does not need actuators to reposition the pellet depending on the paw preference of the animal. For these reasons, desktop devices are simpler, and are therefore less likely to mechanically fail. Finally, as each animal has the same amount of time within the devices on a given training session, the exposure to the device by each animal can be controlled.

Both in-cage devices or benchtop devices would benefit from automation of the assessment of each instance of reach-and-grasp, to avoid the need for experimenters to observe each trial, and to enable scaling up for highly parallel training and assessment. We previously developed in-cage devices which used infrared beam-break sensors to evaluate whether a rodent had reached for a pellet and whether this pellet was dropped before retrieval through a slot (Gadiagellan, 2018); socially housed rats were discriminated using embedded RFID technology. Since developing that prototype, off-the-shelf machine learning algorithms became available which enable key-points (e.g., features of interest such as snout, digits, pellet) to be identified automatically within each single video frame. These “pose estimation” algorithms for animals include DeepLabCut (Mathis et al., 2018) and LEAP (Pereira et al., 2019).

We developed a second prototype using DeepLabCut to assess reach-and-grasp in mice in-cage (Kakanos, 2020). However, these devices were complex in design. To operate, they required 3 stepper motors and their drivers, a custom built distribution board, infrared lights, a camera and a RFID scanner. We found these devices required frequent maintenance owing to their complexity, and scaling their manufacture had its own challenges, limiting the number of devices we could build. We also found that reaching performance and device interaction varied considerably among group-housed mice with access to these devices.

A recently released benchtop device for assessing reach-and-grasp (Bowles et al., 2021) uses the DeepLabCut toolbox and does real-time tracking of mice executing the task; however, this requires expensive locally-installed Graphical Processing Units (GPUs), limiting the number of mice that can be trained simultaneously.

We now describe our third prototype, called ReachingBots. Using a simple dispensing mechanism, computer vision and deep learning, we have automated pellet dispensing and trial classification for highly parallel desktop SPRG training with mice using inexpensive hardware. Rather than detecting the presence of a pellet with DeepLabCut in real-time, we used computationally efficient classical computer vision algorithms that allowed us to execute the automatic re-dispensing of pellets using a low-cost single board computer (Raspberry Pi). A single camera is used to monitor the presence of the pellet in near-real time. Subsequently, each trial is classified automatically as a “success” or “failure” offline using two supervised learning algorithms; a key-point detector (created with a DeepLabCut Convolutional Neural Network) whose output was fed into a Recurrent Neural Network (RNN) for final trial outcome classification. To make the device user friendly, a cross-platform Graphical User Interface was created, requiring no coding expertise to operate devices or analyse videos. We show that ReachingBots assist preclinical stroke research: 30 mice were trained autonomously by 10 devices which automatically quantified long term deficits in reaching performance following unilateral photothrombotic stroke in mice. We discovered that some mice showed deficits whereas others recovered rapidly. We conjecture that the former mice represent reach-and-grasp movements predominantly unilaterally in motor cortex whereas the latter represent reach-and-grasp movements more bilaterally in motor cortex. We propose a strategy for preclinical stroke research that involves training a large set of mice and selecting only those 1) who learn to a predetermined level of success, and 2) who also show large, sustained deficits after unilateral stroke.

## Methods

### Device Overview

ReachingBots consist of two modules: the reaching chamber, and the mechanical, automated dispensing unit (**Figure 1**) designed to deliver single pellets in turn (20mg Dustless Precision Pellets, Purified Rodent diet, Bio-Serve (product number: F0071) in front of each mouse. ReachingBots are referred to in this text as ‘ReachingBot(s)’ or ‘Devices’ interchangeably. The reaching chamber is a clear acrylic box with a lid, with one face that has a centrally positioned 5 mm vertical slot, through which the mouse reaches. The reaching chamber is attached to the dispensing unit by push fit, which allows users to adjust the position of the slot relative to the pellet (positioning the slot to the left, right, or the middle of the pellet) (**Figure 2**). The user can also move the dispensing unit perpendicularly away from the reaching chamber, which then requires the mouse to use its paw rather than its tongue for pellet retrieval. The dispensing module consists of a reservoir, an arm which collects the pellets, a servo motor, which drives the pellet dispensing arm, and a side-on camera facing the slot and pellet obliquely. A schematic of the device can be seen in **Figure 1**, and the dispensing mechanism in **Figure 2**.

**Figure 1:**
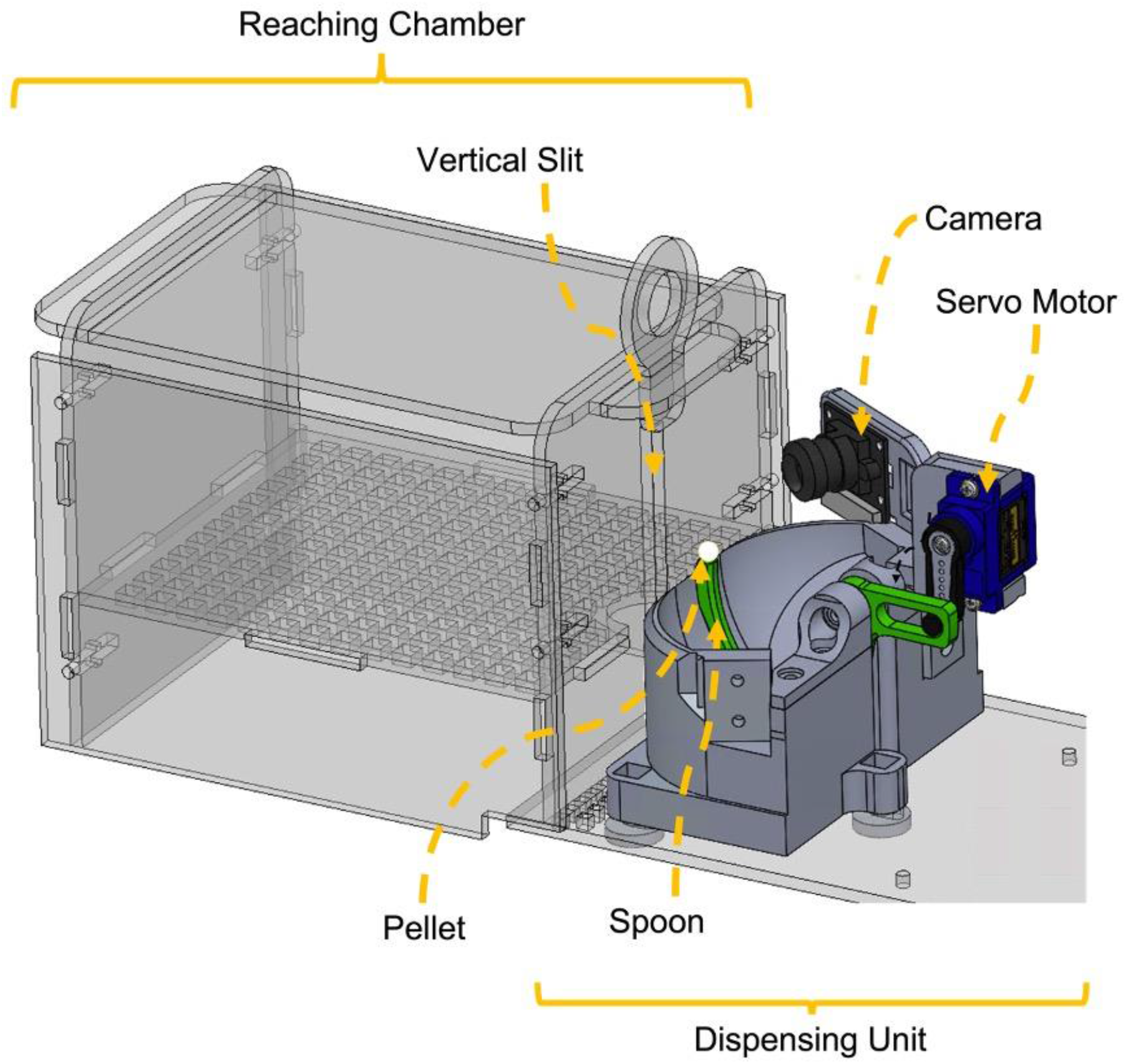
Device overview. **(A)** An orthogonal view of the ReachingBot showing both the Reaching Chamber, where animals are placed, and the Dispensing Unit which shuttles pellets from the reservoir to the vertical slit from which they can be retrieved. The camera can be positioned either to the left or right of the mouse.

**Figure 2:**
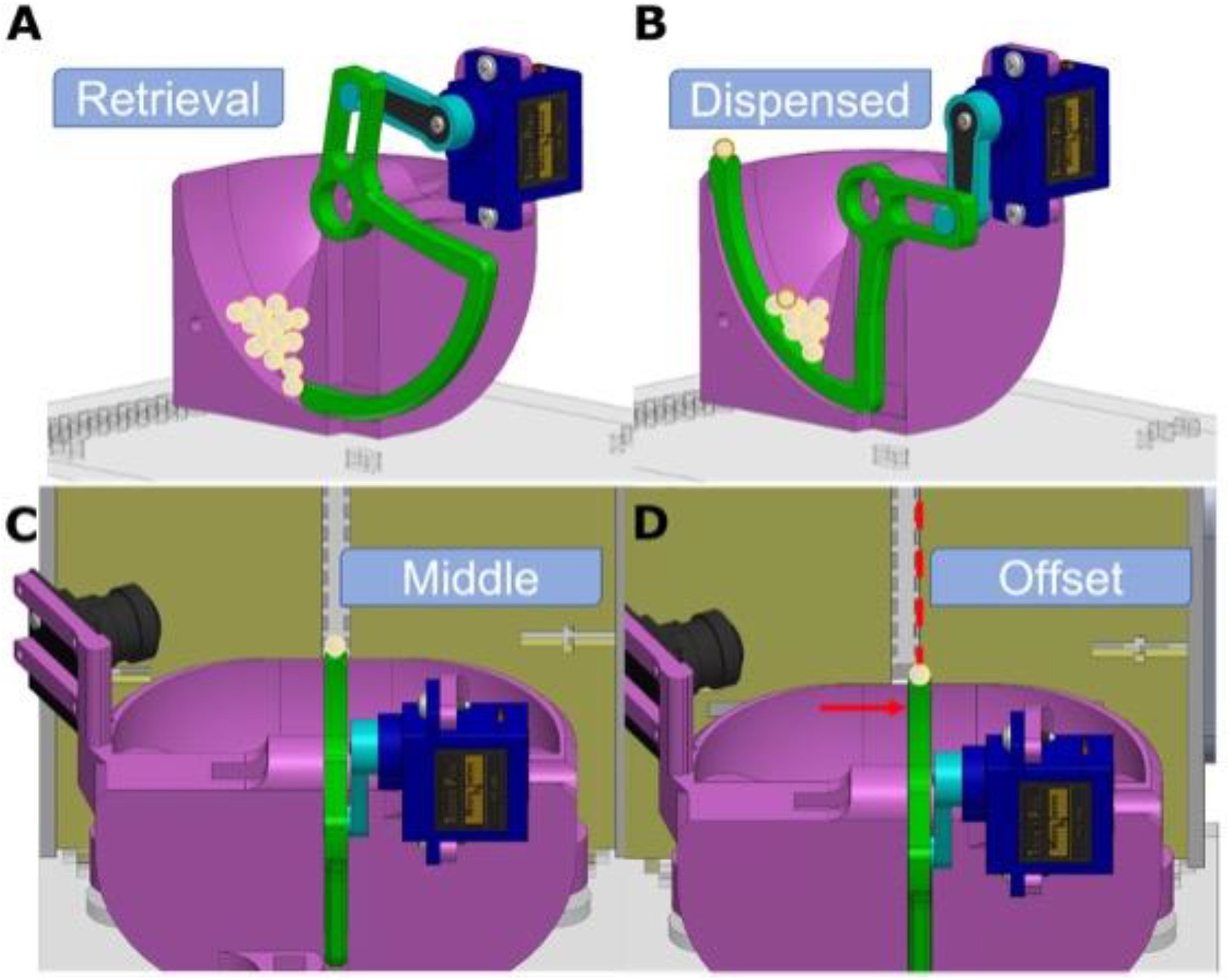
The ReachingBot’s dispensing unit. **(A-D)** The dispensing unit consists of a reservoir (pink), the spoon/arm (green), and the servo-motor couple (turquoise), linked to the servo (blue). The front face plate with the vertical window is denoted yellow. Actuation of the servo motor causes the arm to move from **(A)** a retrieval position to **(B)** a dispensed position. **(C)** During habituation and in the earliest training trials, pellets can be dispensed to the middle of the vertical slot. The distance between the slot and the pellet can be increased progressively (not shown). **(D)** During later training trials, the dispensing unit can be offset to either the left or right of the vertical slot. This allows users to tailor the task to mice with a paw preference, or encourage task participation, respectively.

The distance of the pellet from the slot was configured based on the design described by Farr and Whishaw in 2002 but adapted such that the width of the slot was 5 mm rather than 10 mm (Farr and Whishaw, 2002). The centre point of the pellet is offset from the closest face of the vertical window by 5 mm in the easiest position. In this position, the pellet is central to the vertical window. Mice can use both their paw and tongue to retrieve pellets in this position and is the position used during habituation and early training. This middle position can be offset perpendicularly, so the distance from the pellet to the closest face of the vertical window is 10 mm. In this position mice can only retrieve pellets using either paw, but not their tongue. Finally, a lateral offset can also be applied to either the left or right, by 2.5 mm, meaning the central point of the pellet is aligned with the edge of the vertical window, which we found to be both obstructive enough so the mouse was forced to use the paw contralateral to the pellet offset direction, but not too difficult so that pellets could not be reached at all **Figure 2**. Photographs of the devices with mice reaching for pellets can be seen in **Figure 3**.

**Figure 3:**
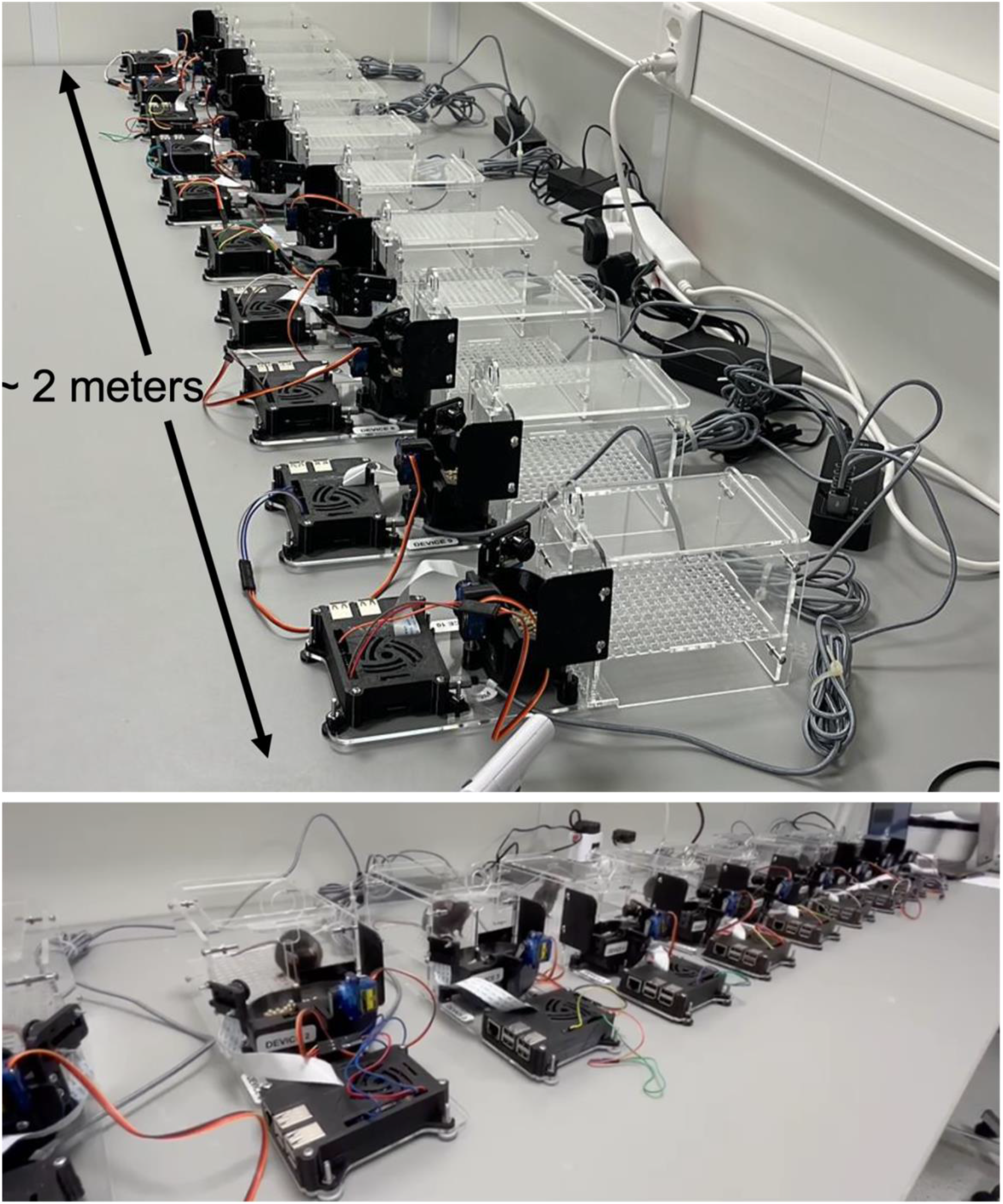
Photograph of 10 devices and mice interacting with ReachingBots. Photographs taken of 10 devices in a row on a workbench occupying approximately 2 meters of space and mice within the reaching chamber, reaching for pellets.

### Software

All the software was written in Python 3, including the firmware (to control the hardware, process the images collected from the camera and classify trials) and the graphical user interface (GUI).

ReachingBots are communicated with, using highly configurable control software: the session length and number of trials can all be configured, for example. Devices are all started simultaneously from one small form factor mini PC (Intel NUC), which acts as a local server to which all of the generated videos are uploaded to for automatic analysis. ReachingBots were configured to automatically present pellets once a pellet was removed from the view of the camera, and to video record the trial with files later sent to the central PC for offline, automatic analysis. The Raspberry Pi handles video acquisition and pellet re-presentation, while the server (having a faster CPU) handles the analysis.

### Component Design, 3D printing and Laser cutting

Hardware was designed using CAD software (SolidWorks 2021), and components to be 3D printed were exported as .STL files. G-code was generated using Ultimaker’s slicing software, Cura. The 3D printed components were fabricated using polylactic acid (PLA) thermoplastic, with a layer thickness of 0.15 mm, a nozzle diameter of 0.4 mm and an infill percentage of 18. The Ultimaker printers were configured to extrude PLA as per the manufacturer’s default settings (210°C temperature). Support material used for components was also in PLA using the ‘tree’ support pattern. The printer used for the manufacture of components was the Ultimaker 2+. Components to be laser cut were rendered to 2D at a scale of 1:1, exported as an Adobe Illustrator (.AI) file, imported into InkScape for line width adjustment to 0.025 mm before being exported as a PDF file (readable by the laser cutter). All of the acrylic components were cut using a 60 watt Epilog laser cutter. The thickness of acrylic (supplier: sheetplastics.co.uk) cut was 3 mm; the parameters used to cut these components were: 10% cutting speed, a frequency of 5000, and at 100% power.

### Onboard Camera

The ReachingBot’s onboard video camera (Raspberry Pi Camera v1, connected to the Raspberry Pi) is positioned to obtain an obliquely-side-on view of the reaching slot. The video camera gathers data for the following tasks: pellet detection, pellet monitoring, and the recording of the reaching attempt. These are all achieved by using the OpenCV library and executed locally on the Raspberry Pi. When active, the camera captures 90 frames per second, with the frame dimensions 640(w) x 480(h) pixels. Captured frames are stored in a cyclical data buffer where up to a maximum of 350 frames could be stored. Additional frames added to the buffer cause the first frame (and oldest) to be removed. Meanwhile, a background function constantly surveys the latest frame for the presence of a pellet (more information in the *Automatic Pellet Dispensing* section). When the pellet is no longer sensed within the frame, frames in the buffer are encoded to a video. In this way, each trial is captured by a single, short video (3.8 seconds, 90 frames per second), avoiding the need to store large amounts of data, and enabling each trial to be reviewed efficiently by the user and/or analysed automatically.

### Automatic pellet dispensing

Pellets are positioned for grasping using a servo motor driving an arm, which lifts an individual pellet from the pellet reservoir. Our previous prototypes involved top-down dispensing (Gadiagellan, 2018), but our current device features a bottom-up dispenser inspired by another group, which was implemented for rats (Torres-Espín et al., 2018) which jams rarely, if ever, and which dispenses a single pellet reliably. Successful pellet dispenses are assessed using a blob detection algorithm which we have previously validated to be a reliable method of pellet detection (Kakanos, 2020). Pellets are brightly coloured within the image frame in contrast to a dark background, meaning they can be detected using OpenCV’s Blob Detection algorithm. We firstly automatically cropped the frame to the region where pellets are placed, applied a Gaussian blur (kernel size of 11 × 11) to the image to reduce noise, inverted pixel values (by subtracting all pixels values from 255), and then applied a the blob detection algorithm.

Blob detection can be optimised to find uniform areas of pixels that are within a given pixel intensity range and that fall within a range of specified surface areas. After empirical testing of these parameters, given the frame size and position of the camera, we found that pellets could be identified robustly by filtering by area so as not to include any blobs which are smaller than 100 pixels. Pellets that have been displaced from the dispensing arm fall outside of the camera’s view and are therefore no longer detected using blob detection, sending a signal to the firmware to encode the video from the frame buffer and collect the next pellet. The blob detection algorithm can be executed on these frames at a rate of 30 Hz.

### Servo motor control

To control the servo motor (Towerpro MG90S), pulse width modulation (PWM) signals were sent to the motor directly from the Raspberry Pi using its PWM peripheral for accurately timed pulses (via a dedicated PWM pin, pin 12). This was done using the gpiozero library. Pulses were sent at 50 Hz, with pulse widths between 0.5 - 2.5 μs depending on the desired arm location.

### Interfacing with and powering devices

Commands and data communications with the devices are handled over USB. All devices could be powered and communicated with simultaneously using a USB power bank (Anker 10 Port 60W Data Hub) connected to one power outlet.

### Cross-platform Graphical User Interface (GUI)

To make interacting with the devices and analysing the videos they generate user-friendly, a cross-platform Graphical User Interface was developed. The app is a standalone desktop application that is locally installed on the computers of users.

#### Device Tab

The app is divided into four tabs. The first is the ‘Device’ tab which is the main segment used to interact with connected devices. Once users have launched the application, sessions begin with the user loading their configuration file [see ‘Setup Tab’ section below for more information] (**Figure 4**) which personalises the interface relevant to the user’s experiments. This is done by navigating to the menu bar and clicking ‘Load configuration file’ under the ‘File’ menu tab.

**Figure 4:**
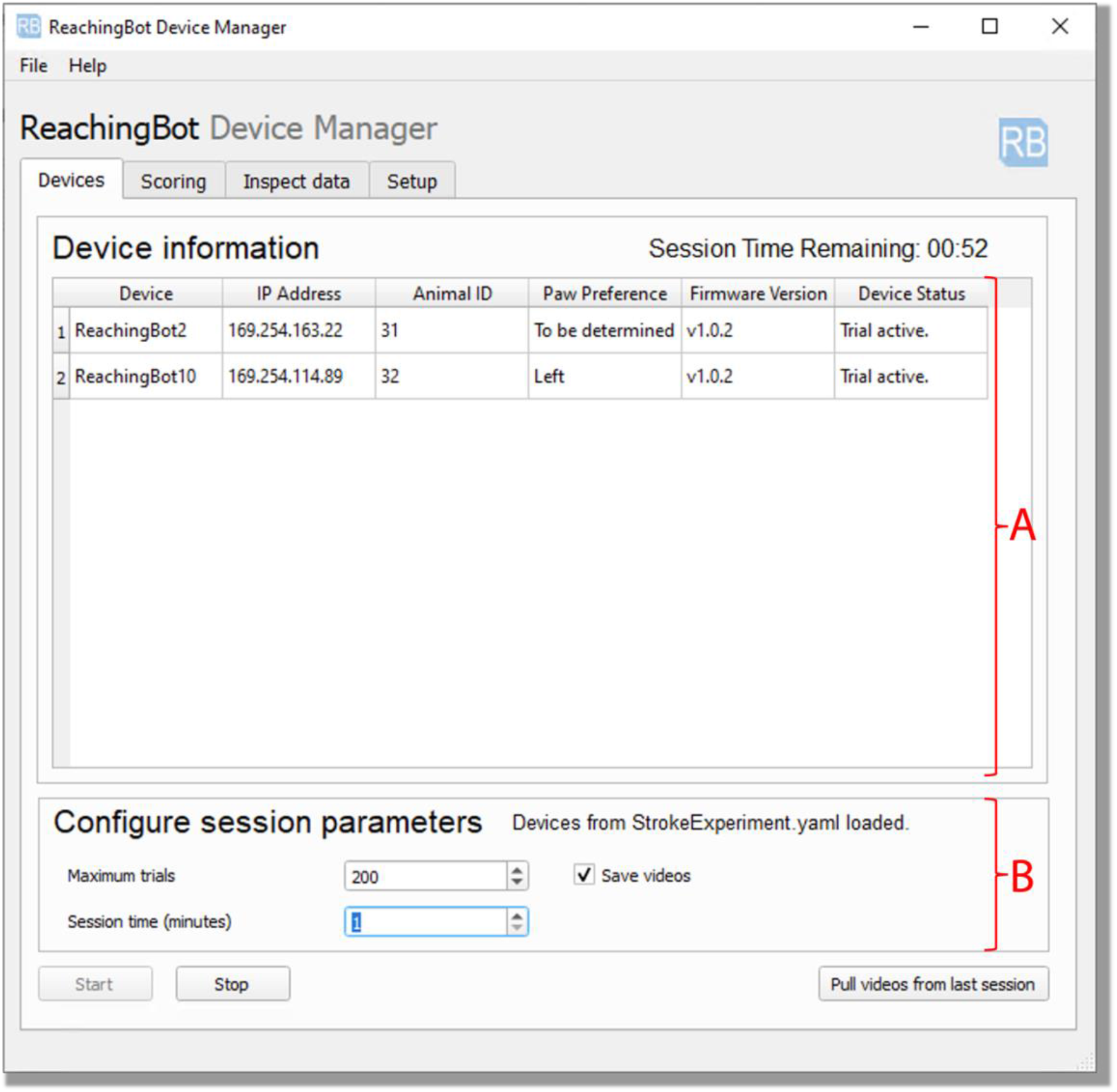
A screenshots of the ‘Device Tab’ from the cross-platform GUI. The “Device” tab allows users to configure the session parameters, including session length, or maximum number of trials, and allowing users to pull the videos from the last session run. Section **A** highlights the table where device information appears, once they are connected to the computer. Section **B** shows where users can change the session parameters before starting devices.

Devices connected to the computer, and configured in the configuration file, will appear in the table below the ‘Device Information’ label in **Figure 4A**. The device name, its assigned local IP address, and its firmware version become visible in the table. Empty fields appear under the Animal ID column, allowing users to enter the identifier for the animal placed in the device within that row. When the animal ID is entered, its paw preference appears in the Paw Preference column, allowing users to configure the ReachingBots so that the chamber is positioned relative to the pellet either to the left or the right, for example. The Device status column displays the last message sent to the host computer by the ReachingBot and signals its state.

Users also have the option to download and install firmware updates to connected devices by clicking on the ‘Update device firmware’ button under the ‘File’ button on the app menu. The ‘Configure session parameters’ section shown in **Figure 4B** allows users to specify the maximum number of trials their animals can take in a given session before pellets stop being delivered, and the maximum amount of time a session will last. In the lowest section of this tab, push buttons can be clicked to either start or stop all devices in parallel. Finally, the ‘Pull videos from last session’ buttons lets users download to the computer all of the videos generated from the session last run, which are generated and stored on the ReachingBots locally. Once videos are uploaded to the computer, they are organised into subfolders according to the date they were generated with the animal ID embedded in the video file name.

#### Scoring Tab

The Scoring Tab (**Figure 5**) lets users classify their videos automatically, leveraging the analysis pipeline outlined in this paper (**Figure 10**). The first panel, highlighted in **Figure 5A**, lets users specify videos for analysis (based on their date of acquisition). The date in this field defaults to today’s date. When the configuration file is selected and loaded the parent directory where videos are stored (as outlined in the Configuration File) is pointed to by default, but users can choose other directories if they wish, by clicking the ‘Select other’ button. In section **Figure 5B**, users can choose which of the subdirectories (i.e. the folder containing the videos generated by a given animal) to run the classification algorithm on, which begins when the ‘Classify Videos’ button is clicked. The benefit of having the choice of which subdirectories to analyse is that the analysis of a given day’s videos can be split across multiple computers, accelerating the process of analysis, especially useful on days when many videos are generated. There are also options to save the coordinates of the key-point detector by checking the ‘Save trajectories’ check box, and also to speed up analysis by having only each nth frame analysed with the neural net. Users can also collate all of the data into one CSV file which will break down the number of reach attempts, successful retrievals and unsuccessful retrievals made by all animals on that given day.

**Figure 5:**
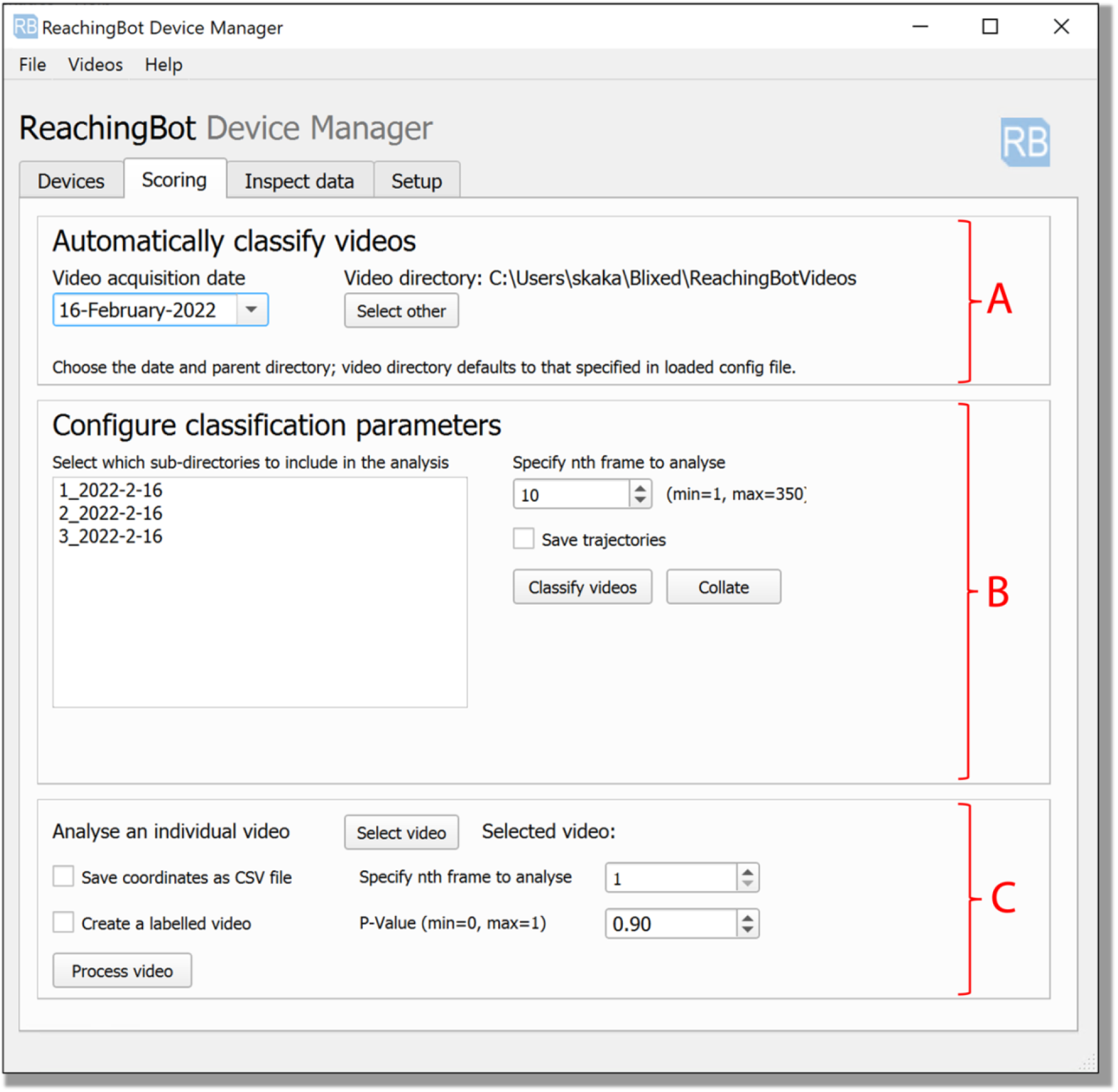
A screenshot of the “Scoring” tab, the interface used for videos to be scored automatically. The Scoring tab is split into three subsections, where in **A,** users can select the date that videos were acquired, in **B** the subdirectories of the parent directory containing videos acquired by a given animal can be selected for analysis, as well as tweak analysis parameters that impact the speed that classifications occur. In **C** users have the option to analyse just one individual video and have the application create a labelled video which overlays the predicted coordinates from the key-point detector over the frames from which they were predicted.

Finally, the section highlighted in **Figure 5C** allows users to generate a labelled individual video so that users can plot the key-point predictions on each frame and inspect mouse paw trajectories. The ‘P-value’ field specified in this section gives users the ability to fine-tune the confidence threshold of DeepLabCut’s output.

##### Inspect data tab

The ‘Inspect data’ tab shown in **Figure 6** gives users a high-level overview of the analysed data generated from the previous scoring tab. Users are given the choice of which subdirectory of data they would like to inspect, and are given metrics such as the number of successful and unsuccessful pellet retrievals made in the session by a given animal (**Figure 6A**). The following section shown in the screenshot in **Figure 6B** features a table showing a session consisting of eight video-recorded trials with the outcome of each trial.

**Figure 6:**
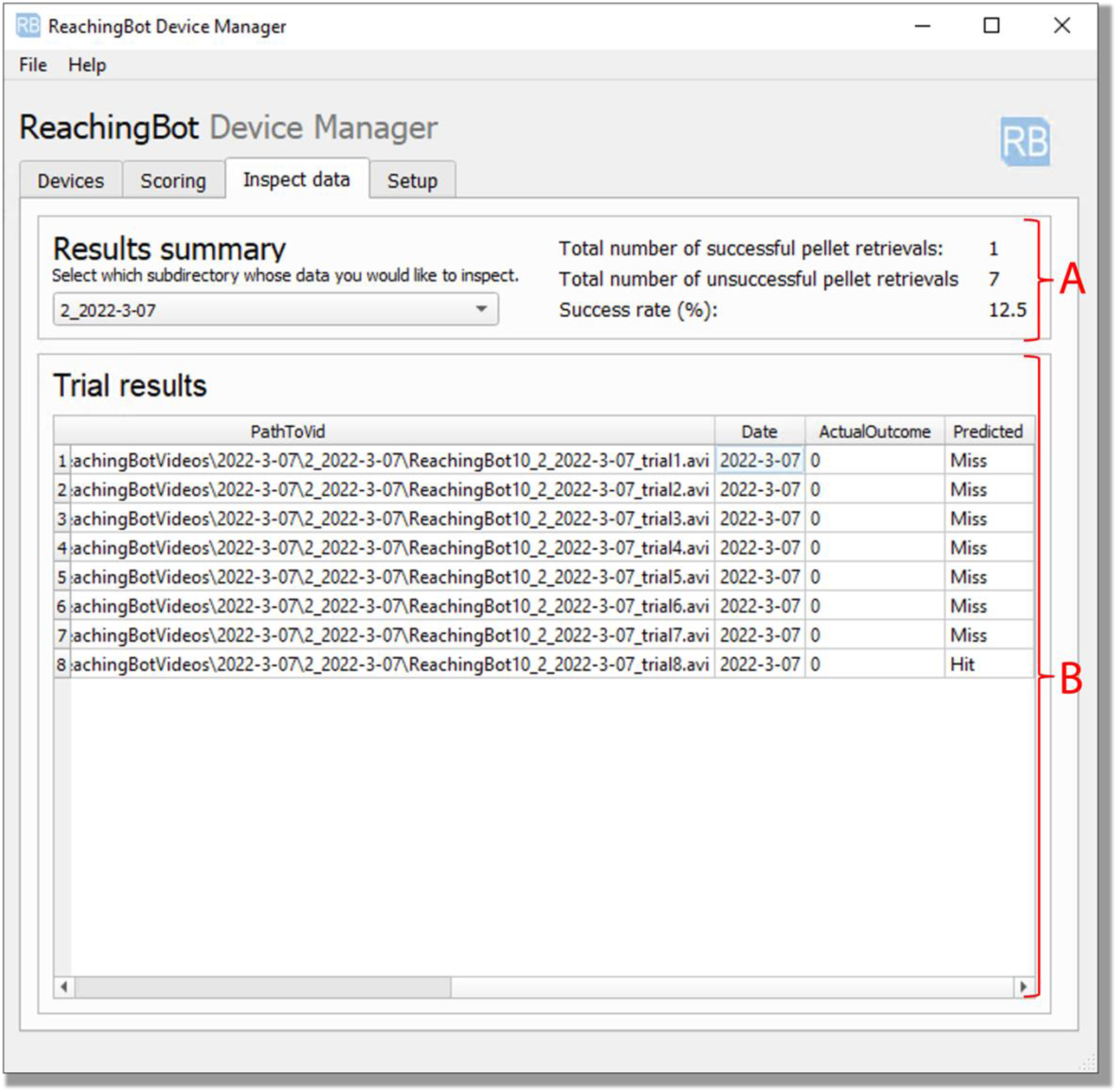
The “Inspect Data” tab allows users to peruse data outcomes quickly, and gives a high-level overview of the classification outcomes. The section outlined in **A** gives users a high level overview of data analysed by the automatic analysis pipeline for a given animal. In **B** a table showing the individual trial outcomes appears.

##### Setup tab

The final tab is the ‘Setup tab’ and is the interface used to configure different experiments. In the project configuration section, shown in **Figure 7**, users can use this interface to create a configuration file, stored as a .YAML file. The first section shown in **Figure 7A**, lets users locate the connected devices on the network and their hostnames, so that the users can either choose to include it in the configuration file, or change its hostname. This can be done by clicking the ‘Start Scan’ button and can be interrupted using the ‘Stop Scan’ button. Here, users can enter the authentication credentials for their devices, and change the hostname of the device from one of the options in the drop-down menu.

**Figure 7.**
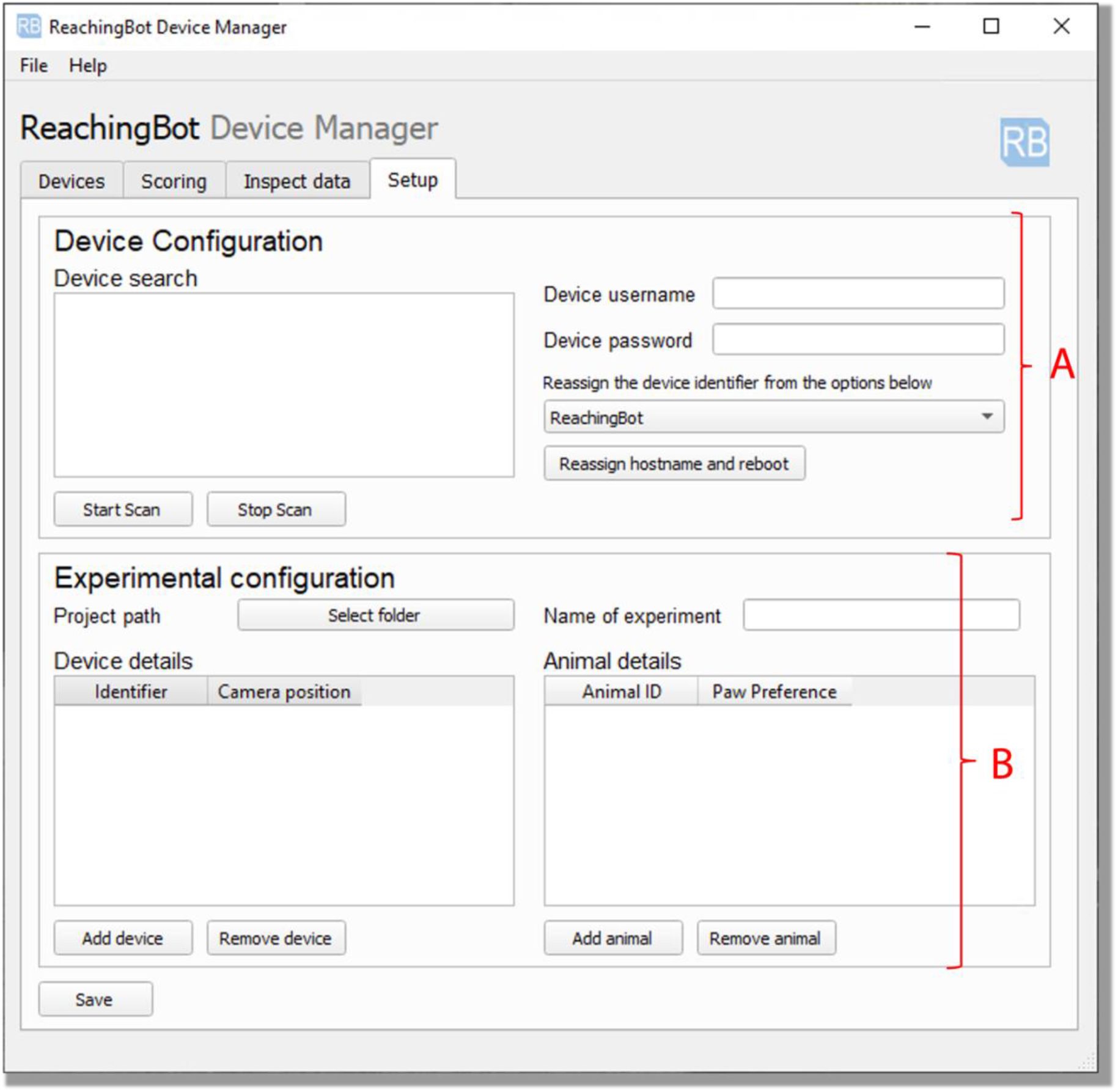
A screen shot of the configuration tab. The configuration tab allows users to create different configuration files that are personalised to their experimental needs, to cater to multiple users, or different experiments. Section **A** allows users to change each device’s unique identifier. Section **B** lets users configure different experiments, by choosing which devices to include, where their cameras are positioned, and by listing the animals used in the study. Users can also enter unique identifier of their animals and their paw preference, which can either be ‘to be determined’, ‘left’, or ‘right’.

In the ‘Experimental configuration’ section, highlighted in section **Figure 7B**, users can specify the folder they would like to store all of the videos and data generated by their ReachingBots, and the name of the experiment. The user can then enter the name of the devices included in this experiment, and the position the camera is in (i.e. ‘Opposite motor’, or ‘Same side as motor’). In the table to the right, users can add which animals are in this particular experiment, their IDs and their paw preference, so that the application can prompt which way to configure the device when they are placed in the reaching chamber. Once these details are entered, this can then be saved by clicking the ‘Saved’ button.

### Animals underwent a shaping period before initiation of training

#### Animals and habituation

To assess the viability of the devices in training mice, a total of 30 C57BL/6 male mice aged 6 weeks old weighing between 25-30 g, obtained from Charles River UK were used. All procedures were in accordance with the Animals (Scientific Procedures) Act of 1986. All protocols involving animals received prior approval by the King’s College London Animal Welfare Ethical Review Board and were authorized by the UK Home Office Project (license number 70/7865, held by Dr. Lawrence Moon). Following arrival, mice were given *ad libitum* access to food and water and were allowed to habituate to the holding rooms for at least 7 days before experiments began. During this period, mice had *ad libitum* access to food and water.

#### Radio Frequency Identification (RFID) Tagging

For RFID tagging, the mice were anaesthetised with isoflurane (5%) in oxygen (1 L/min), and, using a syringe loaded with a glass capsule RFID tag (FDX-B ISO 11784/11785, supplier: Alibaba), tags were injected into the flank subcutaneously before being manoeuvred between the shoulder blades of the mouse. N.b., ReachingBots do not currently automatically identify each mouse based on its implanted RFID tag; rather, we use cost-effective hand-held readers (Halo Scanner by iD Porte) to identify each mouse uniquely.

#### Food restriction

To encourage reach attempts, food chow was removed from the home cages of mice overnight until their shaping, training or assessment session with the ReachingBots finished. After this, mice were allowed *ad libitum* access to food and water for 4 hours in their home cages, which is then removed again until the following session. Mice were weighed daily, and maintained at 90% of their initial bodyweight during food restriction (see **Figure 8A**).

**Figure 8:**
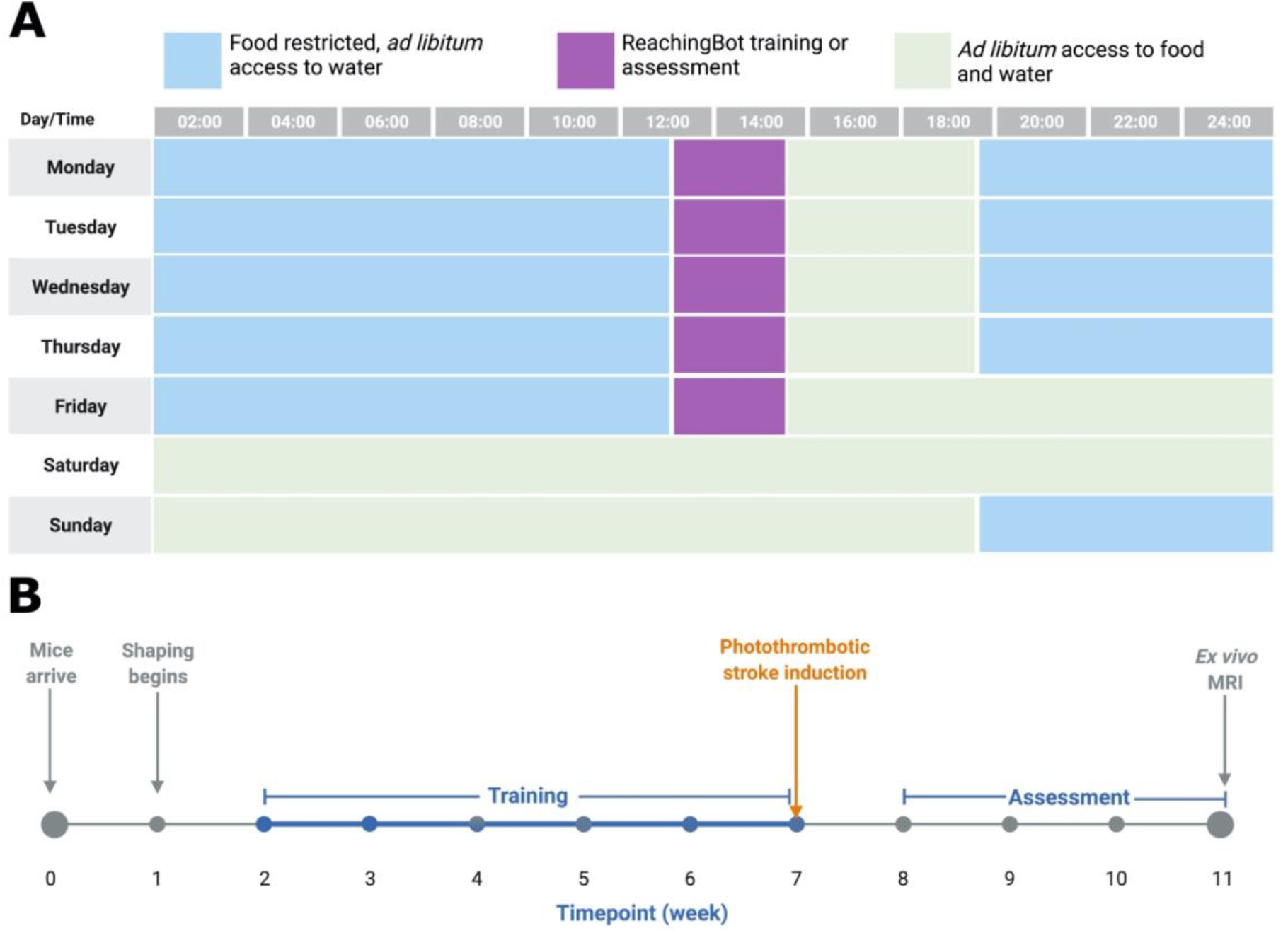
Schematics of the ReachingBot training and assessment schedules during the week, and the overall experimental timeline. The times during the week when mice were either habituated, trained or assessed, along with the food restriction programme is shown in in panel (A). In (B) an experimental timeline is show of the entire study timeline. All schematics were generated using Biorender.

**Figure 9:**
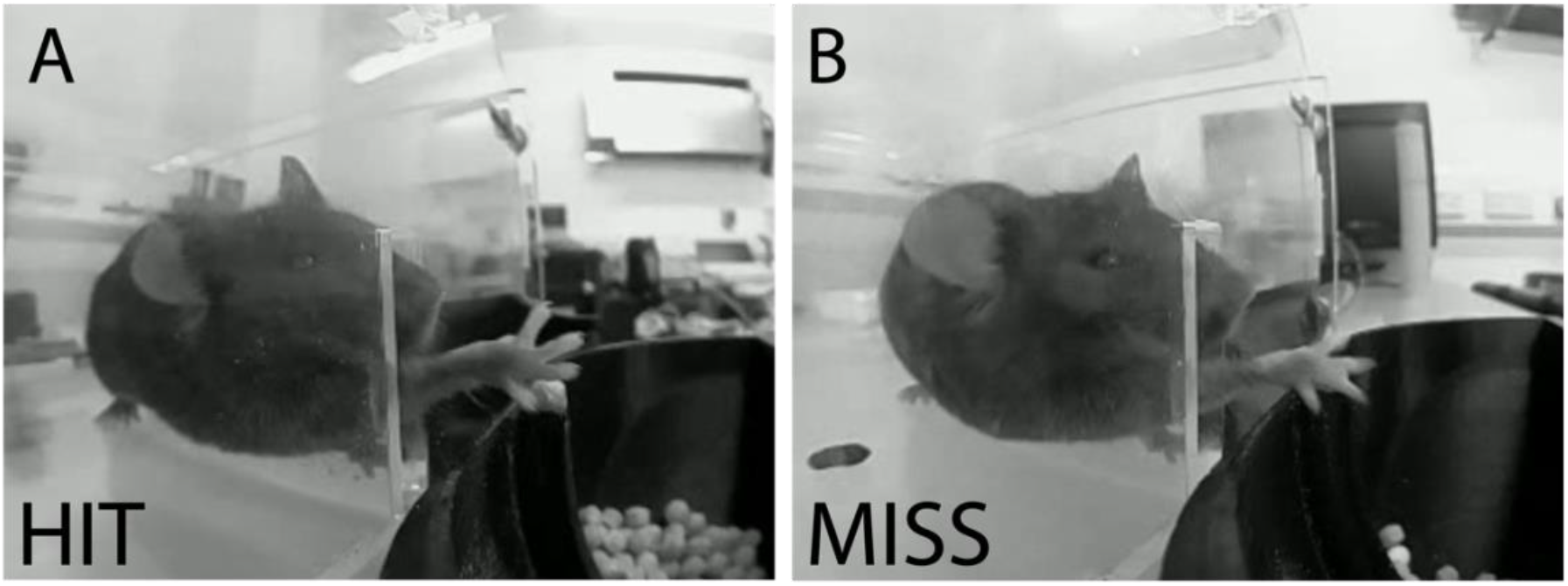
Example screenshots taken directly by ReachingBots of a ‘hit’ (left) and a ‘miss’ (right). Videos taken by the Raspberry Pi camera, at 90 frames per second, with an image size of 320 by 240. A) The video from which this still is taken shows an example of a mouse extending its right paw, flexing its digits around the pellet and grasping it from the first attempt, before retracting its paw, bringing the pellet to its mouth through the vertical slot where it finally uses both paws to support the pellet while the mouse eats the pellet. B) A different mouse extends its paw over the pellet, extends its digits around the paw but drags the pellet over the reservoir’s rim, leading it to fall between the chamber and reservoir. The mouse makes a further reach attempt at the empty arm.

#### Shaping

On each shaping, training and assessment day, mice were placed into the reaching chambers of each device between 11AM and 1PM. The structure of the shaping period is as follows:

**Day 1.** Mice are introduced to the ReachingBot chambers in pairs or trios with the lid locked for 20 minutes. 20mg Pellets are introduced to the floor of the ReachingBot chamber and the dispenser is positioned to deliver pellets just beyond the vertical split so that it can be licked or taken with a paw. The food pellet reward is also added to the home cages of mice overnight.
**Day 2.** Mice are re-introduced to ReachingBots individually as outlined on Day 1. This time, no pellets are placed on the floors of the ReachingBot chambers and can only be retrieved from beyond the vertical slot. The dispenser is still configured to deliver pellets in the easiest-to-reach position; directly in the middle of the vertical slit, with minimal distance from the slit.
**Day 3-5.** The distance of the pellet relative to the vertical slit is now increased, but the pellet remains in the middle of the slit. Mice can now no longer reach for pellets with their tongues and must now use either paw to retrieve the pellet. Users can now look over the video footage generated and determine their preferred paw so that the offset can be applied the following training week. The preferred paw of each mouse was determined by inspecting 20 consecutive videos and assigning the preferred paw as the one used to reach for pellets in more than 55% of trials.

### Training schedule for SPRG

Once shaping was complete and mice could reach for pellets and demonstrate a paw preference, their training, to increase their success rate, could begin. Training sessions began with RFID tags for each mouse being read, then entered into the ReachingBot app (see **Figure 4B**) corresponding to the device the mouse was placed in. Mice were allowed to reach for pellets for 20 minutes with no upper limit on the number of trials per session (this was configured in the section of the app shown in **Figure 4C**). When a pellet was either taken or dropped, a new one was presented to the mouse six seconds later.

### Definitions of trial classifications and performance metrics

Trial outcomes were either a ‘hit’, ‘miss’ or a ‘cheat’ and were used to manually classify trial outcomes of a number of videos to form the training dataset used to make the automatic classifier (**Figure 10**; Supplementary Videos 1 and 2 for example videos).

**Figure 10:**
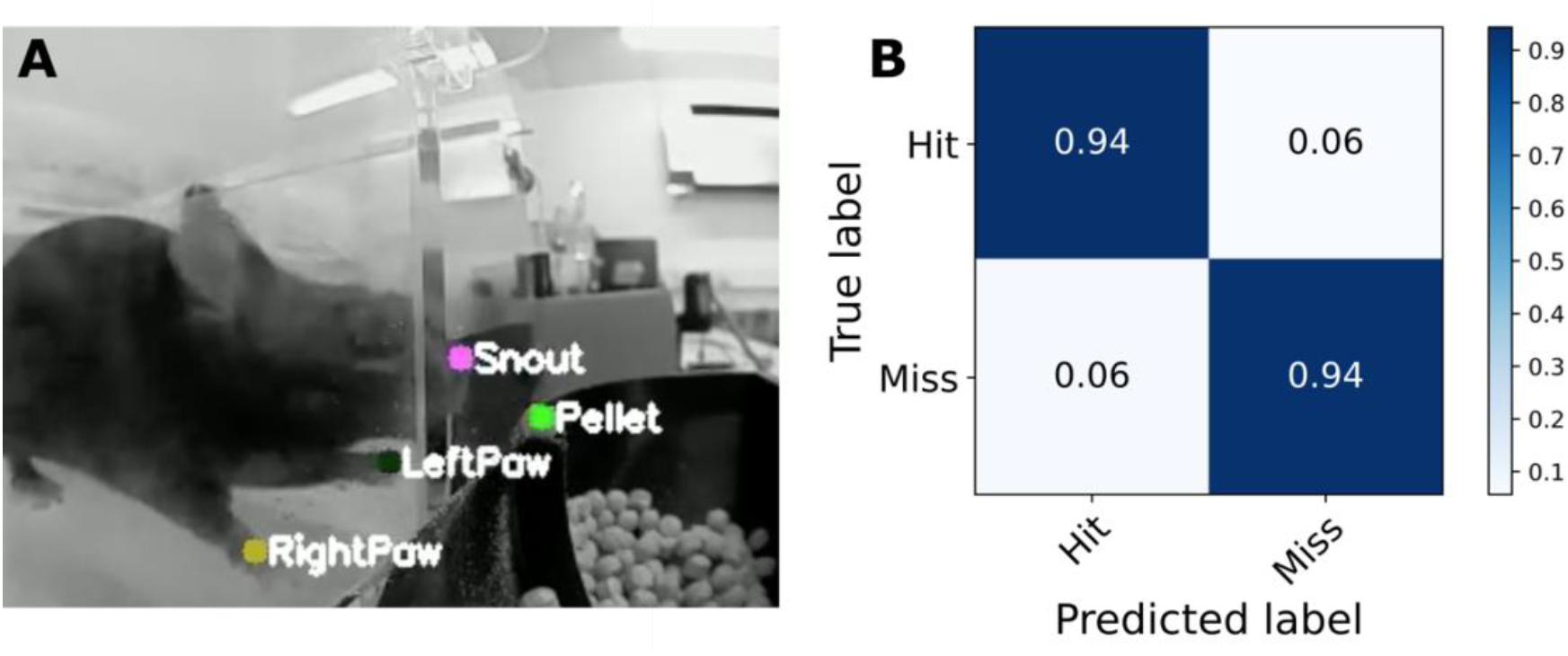
Automatic video classification. **(A)** Four key-points automatically extracted from and overlaid on a video frame: snout, pellet, left paw and right paw. **(B)** A confusion matrix showing the accuracy of our analysis pipeline at classifying video outcomes into ‘Hits’ and ‘Misses’, which both came to 94%.

#### Hit classifications

A trial was classified as a ‘hit’ when a mouse successfully reached and retrieved a pellet, brought it to its mouth without dropping the pellet at any point, irrespective of the number of reach attempts made.

#### Miss classifications

A trials was classified as a miss if the pellet was knocked off the pedestal or dropped at any point of the trial.

#### Cheat classifications

A trial was classified as a cheat if a mouse managed to flick a pellet through the vertical chamber slit, where it was then able to eat the pellet directly from the chamber floor. Instances of this trial were limited by creating a perforated chamber floor such that pellets immediately fell through in the event this happened. When the chamber is positioned 1 cm from the pellet (the furthest configuration) mice are unable to retrieve pellets by licking.

#### Trial

A trial is a rodent’s attempt to retrieve the pellet that results in the pellet being removed from the spoon, whether that be a Hit, Miss, or a Cheat.

#### Success Rate

Success rate percentages for a given day were calculated using the following equation:

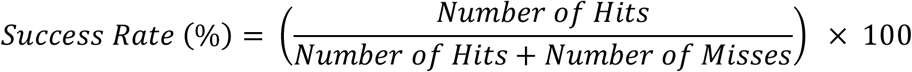

#### Peak Success Rate

To account for intra-week fluctuations in reaching performance, the peak success rate was taken. For a given animal, this was the highest success rate achieved by a mouse on a given day per training week (Monday to Friday).

### Scoring and inclusion criteria

Mice were excluded from reaching performance analysis if they did not successfully reach 40% of their pellets on at least one day throughout the training period; 9 of the 30 mice did not reach this criteria. On a given day, if a mouse did less than 20 trials, the data for that day was not included. The frequency of this was low, with mice typically reaching at least 80 times in a given training session 1 week after the habituation period. Following photothrombotic stroke, a second inclusion criterion was applied such that only mice whose peak reaching performance dropped by 5% or more in the first week following photothrombosis were included into the success rate analysis.

Success rates in **Figure 12** and **Figure 13** were generated by manual classification of the last 20 video trials on a given day.

### Photothrombotic Stroke Surgeries

On the day of the surgery, the mice were weighed, anesthetized by 5% isoflurane, and maintained at 2% isoflurane in an oxygen/air mixture through a nose cone attached to the stereotaxic frame. Fur was shaved above the skull and skin was swabbed using antimicrobial solution (4% chlorhexidine gluconate). Viscotears (Boots, UK) was applied to the eyes of the mice to prevent drying. A midline incision was made along the scalp from above the nose to between the ears, and skin was retracted using weighted clips. The periosteum was removed with the edge of a scalpel. Photothrombosis was confined to a 15 mm^2^ rectangular area (5 × 3 mm) encompassing both forelimb and hindlimb representations. An aluminium stencil was placed to expose a rectangular region of skull located between 0 to 3 mm laterally (hemisphere contralateral to preferred forelimb), and from 2.5 mm anterior to −2.5 mm posterior to Bregma to confine irradiation to the skull overlying the motor cortex. Strokes were unilateral and the side lesioned was contralateral to the preferred paw of mice. We used an optic fibre with a 6 mm aperture attached to a KL 2500 LED light source (Schott) set to produce a light intensity of 20 klux as determined by a light meter (ILM 1335) attached to the fibre optic at a set distance (120 mm) using a 3D printed adapter (Supplementary File 1). The optic fibre was held by a stereotaxic arm, over and in light contact with the aluminium stencil. Once all this was in place, 30 mg/kg of 95% Rose Bengal (Sigma-Aldrich, catalogue number: 330000, diluted in saline) was injected intraperitoneally. Five minutes later the light was turned on for 10 minutes. Saline was applied to the skull following the light activation period and the incision was sutured. Saline (0.5 ml) and pain-relief medication [Carprieve, diluted in saline, (6.5 mg/kg)] were administered subcutaneously post-operatively. Animals were then placed in a warm chamber heated to 33 °C until conscious.

### Automatic trial classification

Automatic trial classification was achieved by using two supervised machine learning algorithms. The first was an automatic key-point detection algorithm (using DeepLabCut, to identify pellet, snout and paws in each video frame) and the second, a recurrent neural network, to predict whether a trial was a ‘hit’ or a ‘miss’).

To generate the training set for the key-point detecting neural network, 640 frames were taken from a range of videos of different C57BL/6 mice executing the reaching task in different ReachingBots on different days, and annotated using DeepLabCut’s graphical user interface. The key-points labelled were the pellet, the snout of the mouse, and the most proximal part of both left and right paws (see **Figure 10A**). Once labelled and the dataset was created, a model was trained by way of transfer learning where the initial weights were that of the ResNet50 model. The model was trained for 500,000 iterations with an Nvidia GeForce RTX 2080 Ti. DeepLabCut version 2.2.0.0 was used throughout.

To generate the training set for the recurrent neural network, 1000 videos which had been manually classified as “hit” or “miss” were used. These 1000 videos were selected from the dataset randomly and included 500 successful pellet retrievals and 500 unsuccessful retrievals.

These 1000 videos were fed through the key-point detection neural network so that the position of each key-point of interest could be predicted for each frame. The output of this DeepLabCut-trained model inference is a matrix of probabilities corresponding to each pixel of each frame, where this probability value is the likelihood of a given feature being present at that particular pixel. These are floating point values that range from 0 to 1, where 1 is a high likelihood the feature is present at that pixel, while 0 means it is not likely. These probability values are then processed to return the coordinate of the pixel with the highest probability a given feature is present, resulting in a CSV file with rows corresponding to each frame of a video, containing the most-likely coordinate of a given key-point within the frame alongside its probability value, or likelihood score, generated by DLC.

The values from these CSV files generated from the aforementioned 1000 videos were transformed into a Numpy array for each video, and were then concatenated into a single matrix. Using scikitlearn’s StandardScalar function, a scalar was created to normalise the DLC-generated data so that the range of values for each column were standardised. Once the scalar was fit, it was saved as a .pickle file, and used to later transform new instances of DLC-generated data for subsequent trial classification. The process of transforming this data ensured that the classifier is trained efficiently.

Finally, the pipeline to train the trial classifier pre-processed the DLC predictions by loading them into a Pandas dataframe, extracting the values only (no index, nor column titles) and stored them in a Numpy matrix of shape 350, 12, where 350 is the number of frames in the video, and 12 is the number of features extracted from each video (i.e. x, y and likelihood values for each predicted key-point), and the scalar transformation was applied to this data. The training set, consisting of 1000 of these matrices, along with the actual outcome of trial (hit, or miss), was used to train the trial classifier. The training set was split into training, testing, and validation subsets with a ratio of 4:1:1.

The dataset contained an equal number of examples which were ‘hits’ or ‘miss’ so as not to introduce biases in the eventual model. In summary, these two neural networks were trained to predict whether or not a given trial was a “hit” or a “miss” **Figure 10B**.

### Ex vivo MRI scanning and analysis

Mice were euthanised using an overdose of pentobarbital and perfusion-fixed with 4% formaldehyde 4 weeks following photothrombosis. The heads of the mice were post-fixed in 4% paraformaldehyde (PFA) for 24 hours before being transferred to phosphate buffer solution.

*Ex vivo* structural MRI images were acquired using a 9.4T horizontal bore Bruker BioSpec 94/20 and a 39-mm volume coil. Heads were scanned four at a time in a custom 3D-printed holder in a 50-ml Falcon tube filled with proton-free fluid (Galden, Solvay). T2 weighted images were acquired using a fast spin-echo sequence: effective echo time = 30 ms, repetition time = 3000 ms, field of view = 25×25×20 mm, acquisition matrix = 250×250×200, isotropic voxel size of 100 μm, scan time = 5 h 44 m.

Lesions appeared heterogenous and hypointense in the T2-weighted images and were difficult to delineate manually (**Figure 11A-C**). We therefore opted to quantify the volume of damaged tissue by subtracting the volume of the upper quadrant of the lesioned hemisphere, (only including spared tissue), from the upper quadrant of the intact hemisphere (**Figure 11F**). Tissue was delineated using a semi-automated segmentation method. This was done with ITK-Snap Version 3.8.0. Segmentation was initiated by first outlining a cuboidal Region of Interest (ROI) with the ‘Active Contour “Snake” tool’. An ROI was defined for the upper quadrants of each hemisphere separately, excluding the cerebellum and olfactory bulb (**Figure 11D-F**). The lower plane of the cuboidal ROI was manually positioned such that it intersected the corpus callosum at its most lateral point (**Figure 14**). Once the ROI was defined, the ‘Speed Image’ was generated using the thresholding algorithm, and ‘seeds’ were placed within the brain quadrant, and allowed to expand until brain tissue within the quadrant was filled. Once both hemisphere quadrants were segmented, their volumes were extracted.

**Figure 11:**
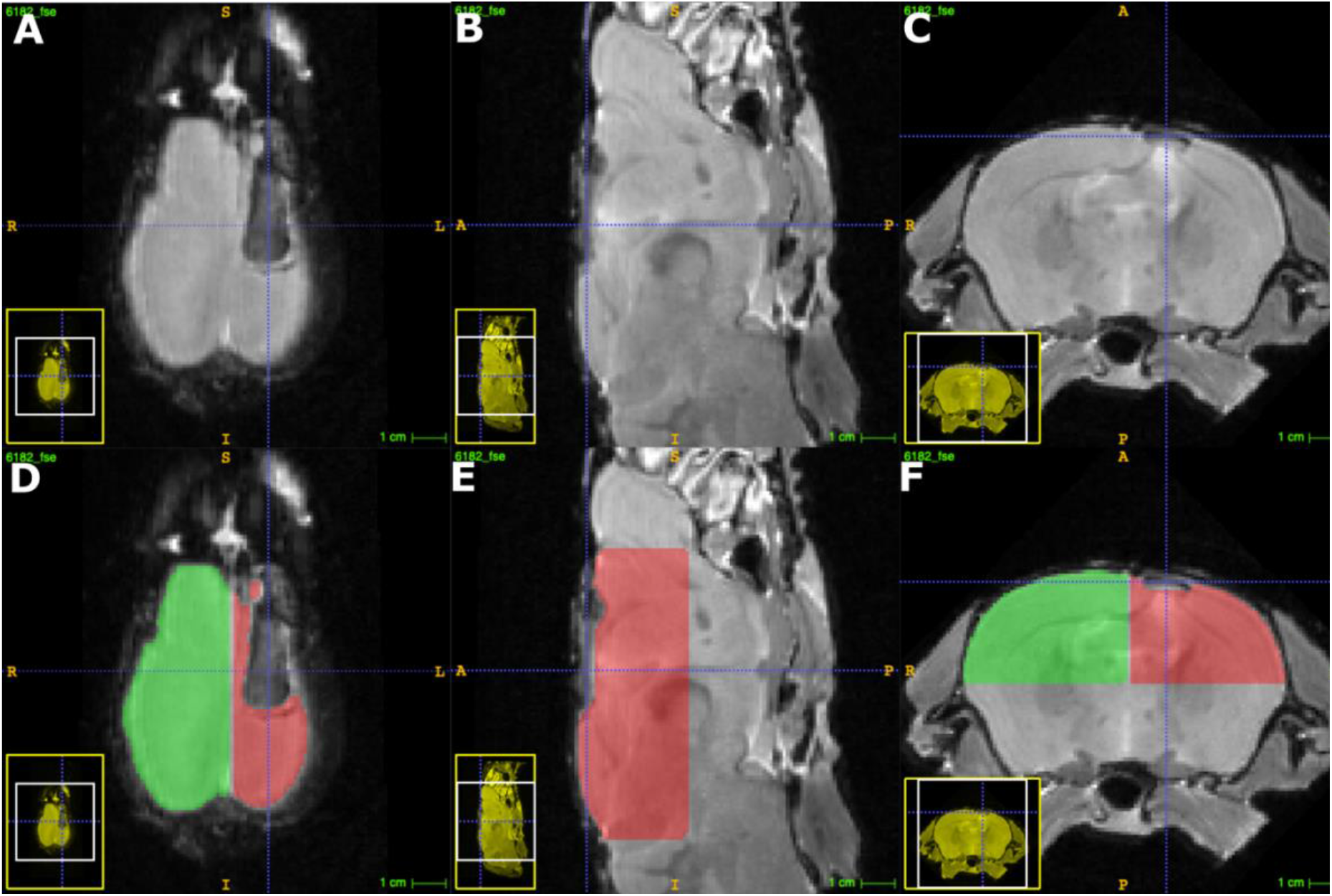
Semi-automated extraction of hemisphere quadrant volumes gives proxy measurement of lesion volume. A representative MRI scan of the mouse brain. A-B show three views of an MRI scan of a mouse brain having undergone unilateral photothrombosis four weeks prior. D-F shows the same sections with the semi-automated segmentation labels overlayed, where green highlights the quadrant in the intact hemisphere, while the quadrant in the lesioned hemisphere is highlighted in red.

### Statistical analyses

Repeated measures ANOVA tests and pairwise comparisons of different timepoints for both daily and weekly reaching numbers and success rates were done using the Pingouin Python package. The threshold for significance, alpha, was set to 0.05. T tests were performed using Python’s Scipy package. The plots were generated using Python and Matplotlib. Graphs and text show mean ± SEM. The threshold for significance (alpha) was 0.05.

## Results

### Trials can be classified automatically using a two-step analysis pipeline

With the hardware and firmware working reliably, and training 30 mice in parallel, thousands of short videos of mice making reaching attempts were being generated each day. Despite each video being confined to the moments before and after the pellet was either dropped or retrieved, scoring them manually would require a large number of hours. We therefore opted to use an automated scoring approach using two supervised machine learning algorithms (see Methods section for more detail). The classification algorithm correctly identified videos as hits or misses in 94% of instances (**Figure 10B**).

### Mice readily interact with ReachingBots and it successfully automates their training and assessment

With food restriction, mice readily interacted with ReachingBots, and, over the course of the training and assessment phase, ~83,000 video trials were recorded, averaging around 2000 trials a day. Using 10 devices, 30 mice were trained unattended, in just under 1 hour and 30 minutes per day, including preparation time. It would take a human doing the equivalent amount of training, over 10 attended hours per day.

Of the 30 mice used in this study, 21 reached a success rate of 40% on at least one occasion. Only mice that reached this success rate in at least one session were included in the groupwise analyses shown next. Over the course of the training period, mice undertook increasing numbers of daily trials within the 20 minute session (**Figure 12A**). With increased exposure to the ReachingBot, the daily success rates of mice also increased (**Figure 12B**). The weekly peak number of trials recorded increased from 79 (+/- 5) after one week of training and peaked at 110 (+/- 6) at the end of week 4 (F(4, 88) = 12.7, p < 0.01, **Figure 12C**). The weekly-peak success rate peaked after 4 weeks of training, with an average weekly-peak success rate of 44.9% (+/- 3%) across 21 mice (F(4, 80) = 8.32, p < 0.01, **Figure 12D**).

**Figure 12:**
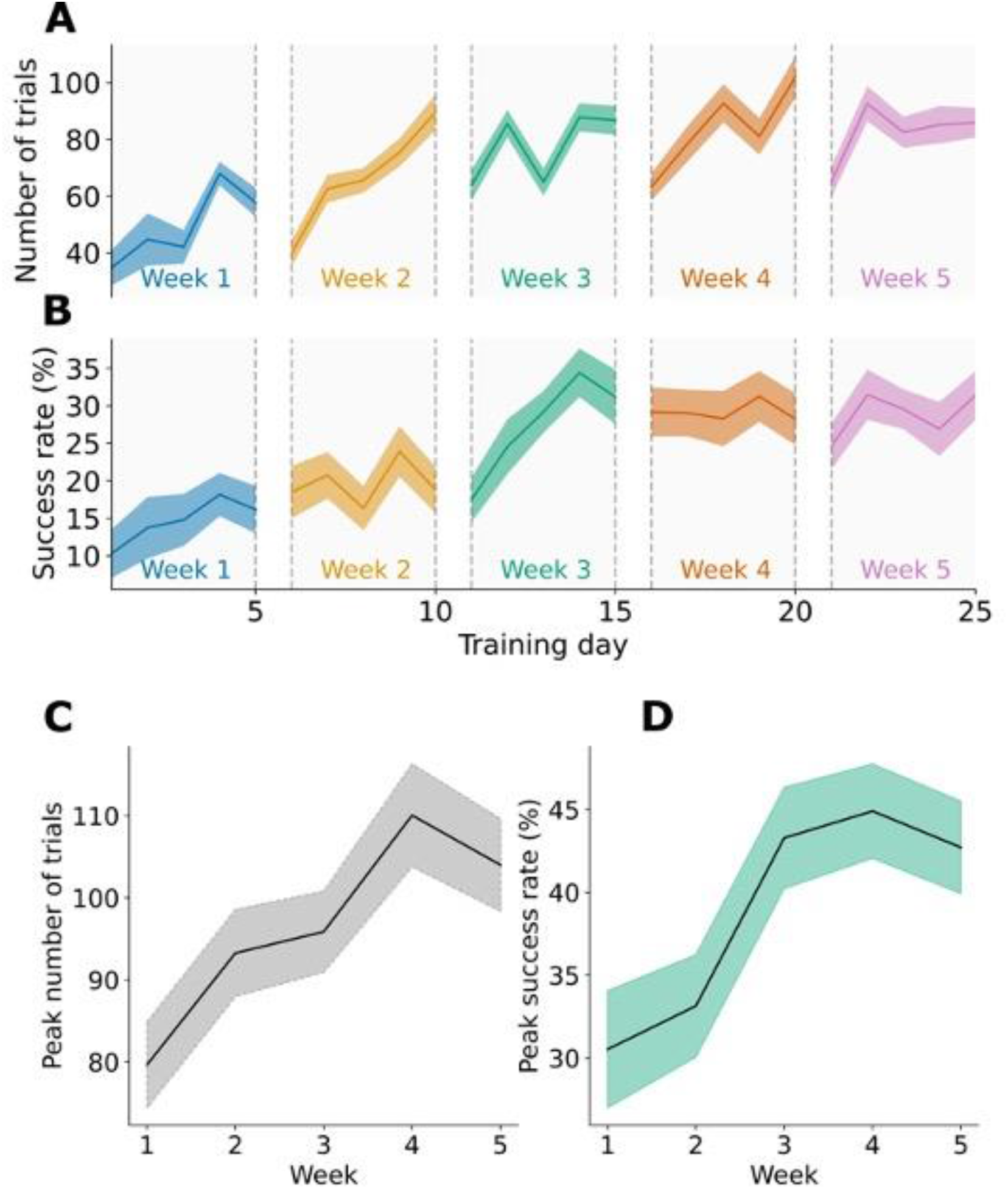
Reaching and Success Rates of mice using ReachingBots increased during the training phase. **(A)** The number of trials performed on the given training day across the 5 week period increased over time. A drop can be seen between each break for the weekend. **(B)** During the training phase, mice improved in reaching performance, with their average success rate increasing over time. **(C) Mice undertook more trials; th**eir groupwise average weekly peak maximum trial number increased over the course of the training period **(D)** The groupwise average weekly peak success rate of mice increased over time. All error bars for each plot are standard error of mean (SEM). All subpanels were generated from data collected from mice that met the inclusion criteria, n=21.

### Photothrombosis of the motor cortex resulted in heterogenous reaching outcomes

Following photothrombosis, the motivation for mice to keep reaching for pellets was sustained, with the number of reaching attempts at week 7 not differing to their level pre-injury (**Figure 13A, C**). Only a subset of mice met the inclusion criteria after stroke (9 out of 21), of which, group mean peak success rate dropped from 52% (+/- 4) to 25% (± 3%) one week post stroke (two sided paired t-test, p<0.01; **Figure 13B, D**) with the deficit sustaining over the 4 week post stroke assessment phase.

**Figure 13:**
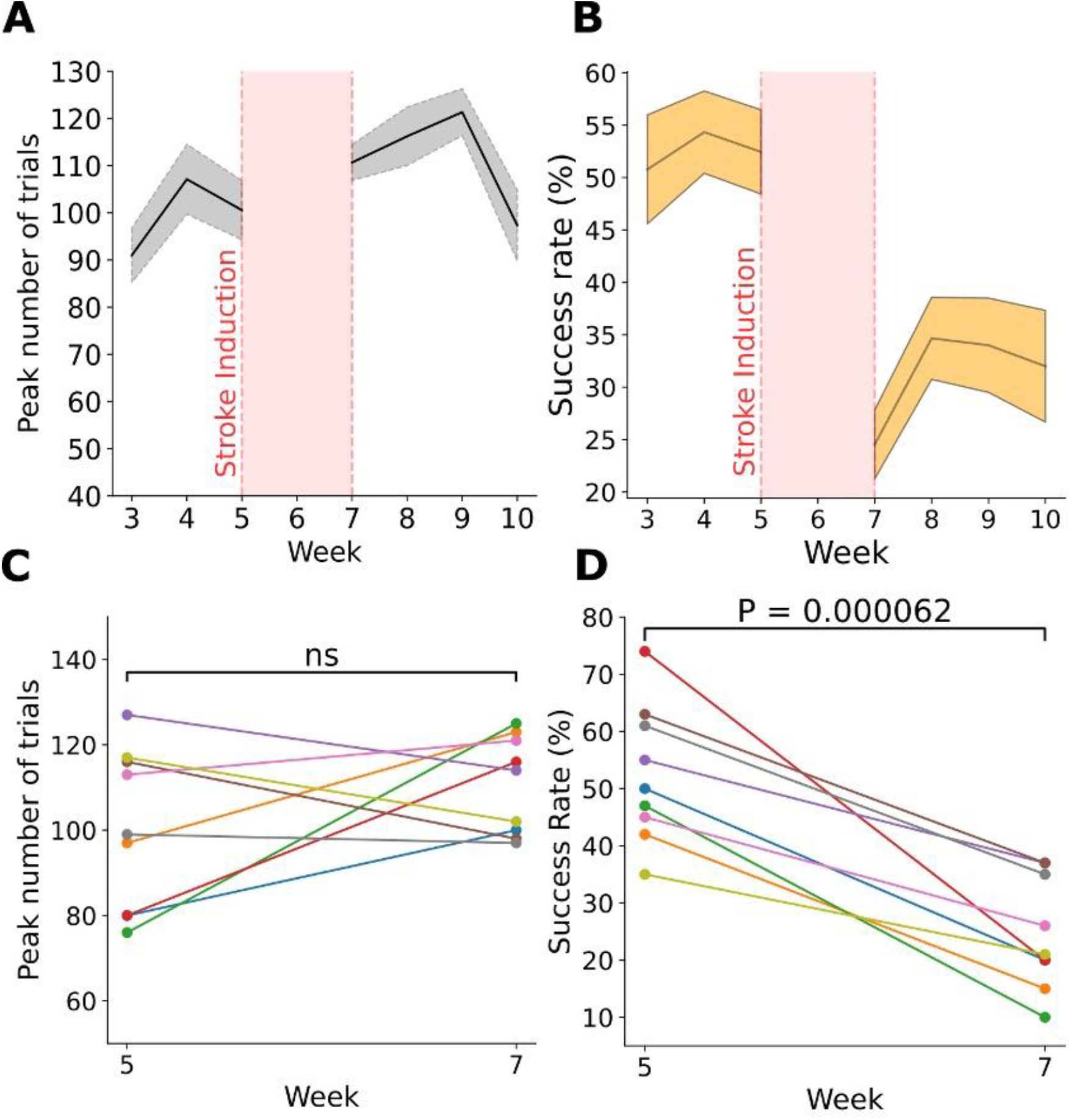
ReachingBots detect drop in reaching performance following photothrombotic stroke. **(A)** Average peak number of trials for the given training week among the nine mice (n=9) who met the inclusion criteria following photothrombotic stroke **(B)** The average peak success rates of mice (n=9) who met the inclusion criteria before and after photothrombotic stroke. **(C)** and **(D)** show the paired number of trials and peak success rates at the baseline (week 5) and at 1 week post injury for the mice who met the inclusion criteria; the colours for each plot are matched to the same mouse.

### Structural MRI revealed that lesion size was not correlated with reaching outcome following stroke

We next wanted to establish whether the heterogeneity of reaching outcomes could be explained by differences in lesion volumes 4 weeks following photothrombosis. To do this, *ex vivo* T2-weighted structural MR images were acquired. A representative lesion can be seen in **Figure 10** and in **Figure 14A** denoted by the red arrow. Photothrombosis caused unilateral loss of motor cortex parallel to the midline in a shape that was approximately cuboid and measuring 5 mm in length. The corpus callosum and underlying subcortical grey matter were displaced dorsally into the zone where the motor cortex had previously existed. Lesions were quantified by subtracting the volume of spared tissue within the lesioned upper brain quadrant from that of the contralateral quadrant. A 3D reconstruction of the extracted segments revealed a crater-like dent at the site of injury (**Figure 14B**). The average lesioned quadrant was 95.3% the volume of the intact quadrant (Paired t-test, P < 0.001) (**Figure 14C**). A correlation analysis of the lesion sizes of all animals and their behavioural outcomes showed that the lesion volume did not predict the change in reach-and-grasp peak success rate (Pearson’s Correlation Coefficient= 0.104, P-value = 0.65, n = 21).

**Figure 14:**
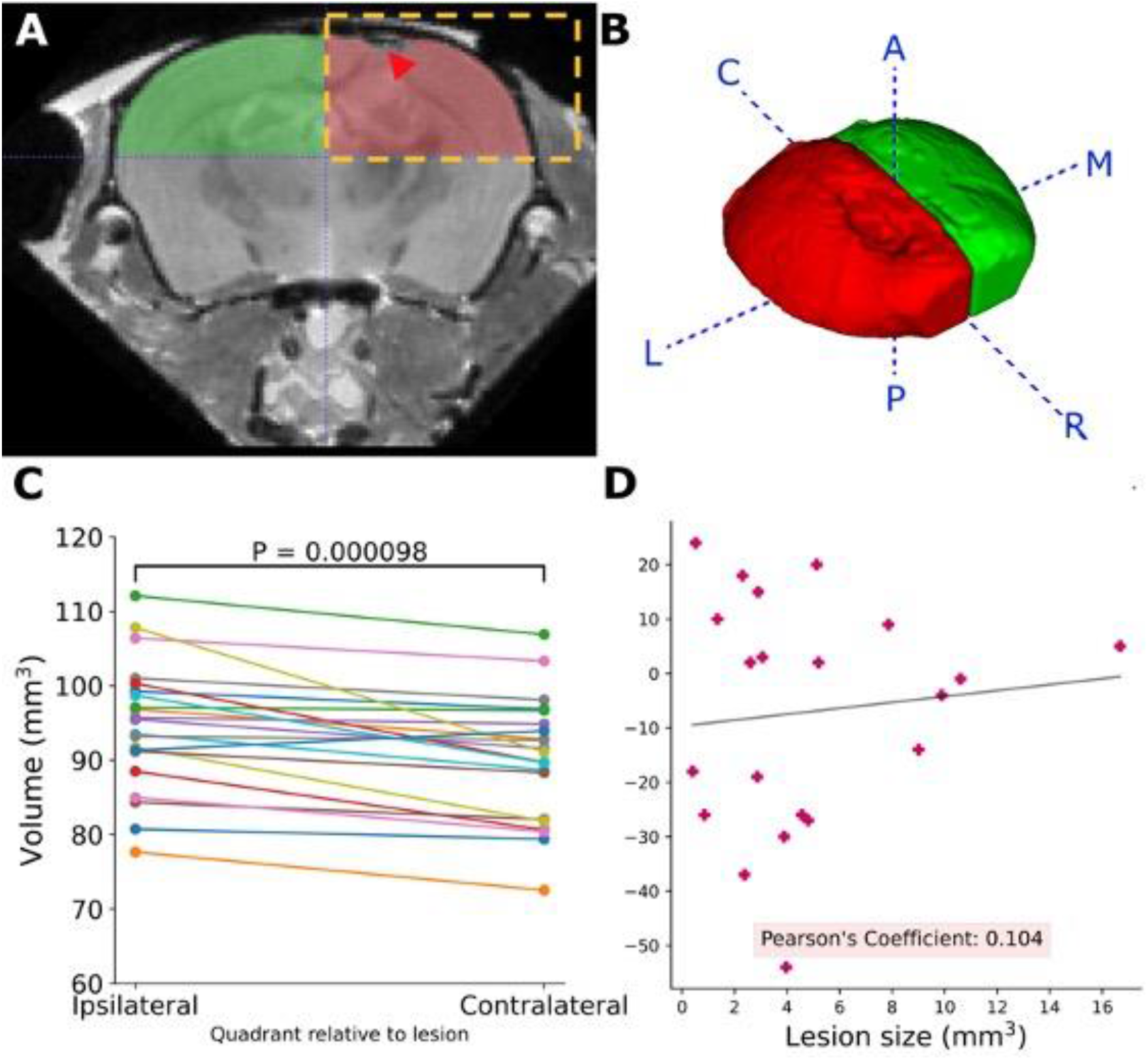
Structural MRI reveals that lesion volume was not correlated with reaching outcome following stroke. **(A)** A coronal slice of a mouse brain having undergone ex vivo structural MRI. A representative lesion can be seen in the right hemisphere, and is denoted by the red arrow. The region of interested (ROI) used in the semi-automated segmentation process can be seen outlined by the dashed rectangle. The corpus callosum can also be seen displaced into the area where the motor cortex previously existed, below the lesion site (red arrow). **(B)** A representative 3D reconstruction of the segmented, post-stroke upper hemisphere quadrants (same brain from which the image in (A) was taken). **(C)** Quantification of lesioned hemisphere quadrant volumes. The lesioned hemisphere was lower in volume than that of the intact hemisphere **(D)** A scatter plot showing the percentage change in success rates of animals before and after stroke against the lesion volume.

## Discussion

### ReachingBots can be used reliably in preclinical studies and increases researcher productivity

Ischaemic stroke is a leading cause of disability worldwide and new therapies are desperately needed. Assays to interrogate movement disorders in animals remain an invaluable tool for the development of novel clinical drug therapies, and the advancement of low cost electronics and robotics mean that these assays can now increasingly be automated. In this paper we describe a novel device that automates the single pellet reaching task for mice in a way that is scalable, low cost, reliable, and easy to use. Currently, benchtop devices that exist to automate the single pellet reaching task in rodents either require that users manually press buttons (Fenrich et al., 2016) to re-dispense a pellet for retrieval, manually classify trial outcomes, or require expensive compute-hardware to run (Bowles et al., 2021). The device described in this paper is unique and an advancement to the now-available technologies in that pellet presentation is handled automatically, trial classification is handled by the newly developed cross-platform desktop application, and all of this can be done without the use of expensive GPUs.

### ReachingBots automatically train and assess mice on the single pellet reaching task

We found that with food restriction, mice readily interacted with ReachingBots, and, in a 20 minute session, attempted to retrieve approximately 80 pellets on average. Collectively, the 30 mice in this study performed approximately 83,000 trials across 10 devices. Human operators will typically present 30 pellets to animals over the course of 20 minutes in the manual version of this task (Chen et al., 2014, Tang et al., 2018), which requires their full attendance. ReachingBots successfully automated the training of 30 mice, where the equivalent time required to do this task would be over 10 hours of work per day, rather than 1 hour and 30 minutes. Given that it typically takes a human operator 20 minutes to manually train or assess an animal doing SPRG, ReachingBots provide a time saving and scalable parallelised training paradigm. Increasingly, dose or intensity of rehabilitation is being shown to be an important predictor of post-stroke and spinal cord injury recovery, with some preclinical drug trials requiring animals to do 100 trials in a session (Wahl et al., 2014). With ReachingBots these studies become increasingly viable.

### ReachingBots train mice in parallel and the majority of participating mice reach success rates of upwards of 40%

Of the 30 mice that received daily training with ReachingBots, 21 reached a pre-defined criteria of a 40% success rate at least once during training. It is possible that with more training, more mice could reach this criteria, as a subset of mice began reaching more successfully later in the training phase.

We find that an efficient way to run a preclinical trial (e.g., to evaluate a new intervention for stroke) using ReachingBots is to obtain a larger number of mice than is required for the actual experiment (e.g., 50% more than is needed). These mice can be habituated and trained on the ReachingBot, and then the user retains for the experiment only those mice that learn to reach-and-grasp above a pre-determined criterion (e.g., a success rate exceeding 40%). We do not recommend that users strive to train recalcitrant mice; rather, those mice can be allocated to other experiments not involving reach-and-grasp (or assessed on other outcomes such as walking on a horizontal ladder). In this way, highly parallel training using ReachingBots makes it possible to generate large cohorts of mice that reach-and-grasp comparably well, which in turn improves the power of an experiment, for example, to detect differences between groups.

### ReachingBots highlight the heterogeneity of reaching outcomes following photothrombosis

To test whether ReachingBots could reliably detect a deficit after injury, we opted to use the photothrombotic stroke model, confined to the motor cortex contralateral to the animal’s preferred paw, such that cortical neurons, including corticospinal tract neurons, were ablated unilaterally. We found that just 9 of the 21 mice who reached criteria during the training phase dropped 1 week after stroke to below a success rate of 5% of the pre-injury performance level.

To begin trying to explain the heterogeneity in reaching success after injury, the brains of mice underwent *ex vivo* structural MR imaging. We speculated that there might be variability in the sizes of lesion caused by photothrombosis and that this could explain the observed behavioural outcomes: namely, that only a subset of mice had sustained deficits in reaching performance. While there was variability in the degree of lost cortical tissue following stroke, gross lesion volume did not correlate with the change in reaching outcome. This could be for a number of reasons: for example, gross lesion volume might be less important than its position, and that correlations might be drawn from the coordinates of the lesion relative to bregma, and the behavioural outcome. However, our lesions were found in a consistent location relative to Bregma. Further, because mice were consistently lesioned contralateral to their preferred paw, it might be that a subset of mice have representation of reach-and-grasp programs more bilaterally, and that some might have one that is more unilateral. Further analysis is required to relate the two.

### Machine learning can be used to predict trial outcome automatically, at scale

To process the large number of videos generated by ReachingBots, we sought to automate video classification using two supervised machine learning algorithms. The first, a key-point detector, created using DeepLabCut, extracts key-points from each frame of the video; these are the snout, the pellet, and each paw. The second supervised machine learning algorithm is a binary classifier, that takes the key-points generated from the first algorithm and classifies whether the video is likely to be a ‘hit’ or a ‘miss’. As well as having a state-of-the-art accuracy score of 94%, running these algorithms to classify each video takes just seconds on a CPU (and does not need a GPU), processing up to 50 frames per second for the videos generated by ReachingBots. This classification pipeline is integrated within the cross-platform desktop application described in this paper, meaning no coding-experience is required by the user to implement it. An advantage of inference needing only a CPU, is that the analysis of videos generated by ReachingBots can be done easily across multiple computers running in parallel. For example, in the experiment described in this paper, we synchronised the computer directory containing the video generated by the ReachingBots on Google Drive, allowing them to be accessed and analysed from any connected PCs, and analysed the videos across multiple desktop PCs.

The ReachingBot joins an increasing number of devices aimed at automating preclinical motor behavioural assays for the acceleration of drug discovery (e.g., for stroke and spinal cord injury). ReachingBots and the ReachingBot Device Manager software package solve many of the key challenges with automating the single pellet reaching task. For example, our system automatically handles both the scoring of trials and the re-dispensing of pellets. Our system allowed this to be done at scale, meaning many mice can be trained in parallel. The use of a camera and computer vision algorithms is one of ReachingBot’s main advantage over other similar devices; the camera is used as a sensor for pellets and the algorithm that handles this runs on the ReachingBot itself, rather than a GPU on a PC, drastically reducing the per unit cost. ReachingBots record every trial made by each mouse and stores them as a compact, short video, allowing users to either manually validate the outcome of the trials themselves, or run their own bespoke analysis on the video. To our knowledge, the cross-platform application described in this paper is the first of its kind, and greatly simplifies the setup and device management process for users.

The device described in this paper has few moving parts and is relatively simple to scale; for this reason, its uptake and usage among many research centres is very achievable, and could standardise the single pellet reaching task in mice, and make preclinical studies using this behavioural outcome far more reproducible.

## Acknowledgments

The research leading to these results has received funding from the European Research Council under the European Union’s Seventh Framework Programme (FP/2007-2013) / ERC Grant Agreement n. 309731 (a “Starter-Consolidator” grant) as well as a “Proof of Concept” grant from the ERC (“Miniature Robots”, n. 713410). This device was developed as part of a grant funded by the Medical Research Council to LDFM (MR/S026053/1). Earlier versions of this device were developed with funding to LDFM from Brain Research UK 201617-04) and a PhD studentship from the Rosetrees Trust (CM533).

